# Inferring the demographic history of aye-ayes (*Daubentonia madagascariensis*) from high-quality, whole-genome, population-level data

**DOI:** 10.1101/2024.11.08.622659

**Authors:** John W. Terbot, Vivak Soni, Cyril J. Versoza, Susanne P. Pfeifer, Jeffrey D. Jensen

## Abstract

The nocturnal aye-aye, *Daubentonia madagascariensis*, is one of the most elusive lemurs on the island of Madagascar. The timing of its activity and arboreal lifestyle has generally made it difficult to obtain accurate assessments of population size using traditional census methods. Therefore, alternative estimates provided by population genetic inference are essential for yielding much needed information for conservation measures and for enabling ecological and evolutionary studies of this species. Here, we utilize genomic data from 17 unrelated individuals — including 5 newly sequenced, high-coverage genomes — to estimate this history. Essential to this estimation are recently published annotations of the aye-aye genome which allow for variation at putatively neutral genomic regions to be included in the estimation procedures, and regions subject to selective constraints, or in linkage to such sites, to be excluded owing to the biasing effects of selection on demographic inference. By comparing a variety of demographic estimation tools to develop a well-supported model of population history, we find strong support for the species to consist of two demes, separating northern Madagascar from the rest of the island. Additionally, we find that the aye-aye has experienced two severe reductions in population size. The first occurred rapidly, approximately 3,000 to 5,000 years ago, and likely corresponded with the arrival of humans to Madagascar. The second occurred over the past few decades and is likely related to substantial habitat loss, suggesting that the species is still undergoing population decline and remains at great risk for extinction.

## INTRODUCTION

The aye-aye, *Daubentonia madagascariensis*, (Figure 1a) is one of the most unique of over one hundred species of lemur endemic to the island of Madagascar (IUCN 2024). Initially described in the late 1780s, it is an elusive species in part due to its nocturnal lifestyle, and was believed to be extinct until being rediscovered in the 1950s (Gotch 1995; Kay and Kirk 2000). As the sole extant member of the Daubentoniidae family, the aye-aye occupies a unique position as an early diverging lineage of the lemuriform clade (alongside Cheirogaledia, Indriidae, Lepilemuridae, and Lemuridae) having branched roughly 54.9 to 74.7 million years ago (Yoder 1997; Horvath *et al*. 2008; Perelman *et al*. 2011; Mclain *et al*. 2012).

**Figure 1.**
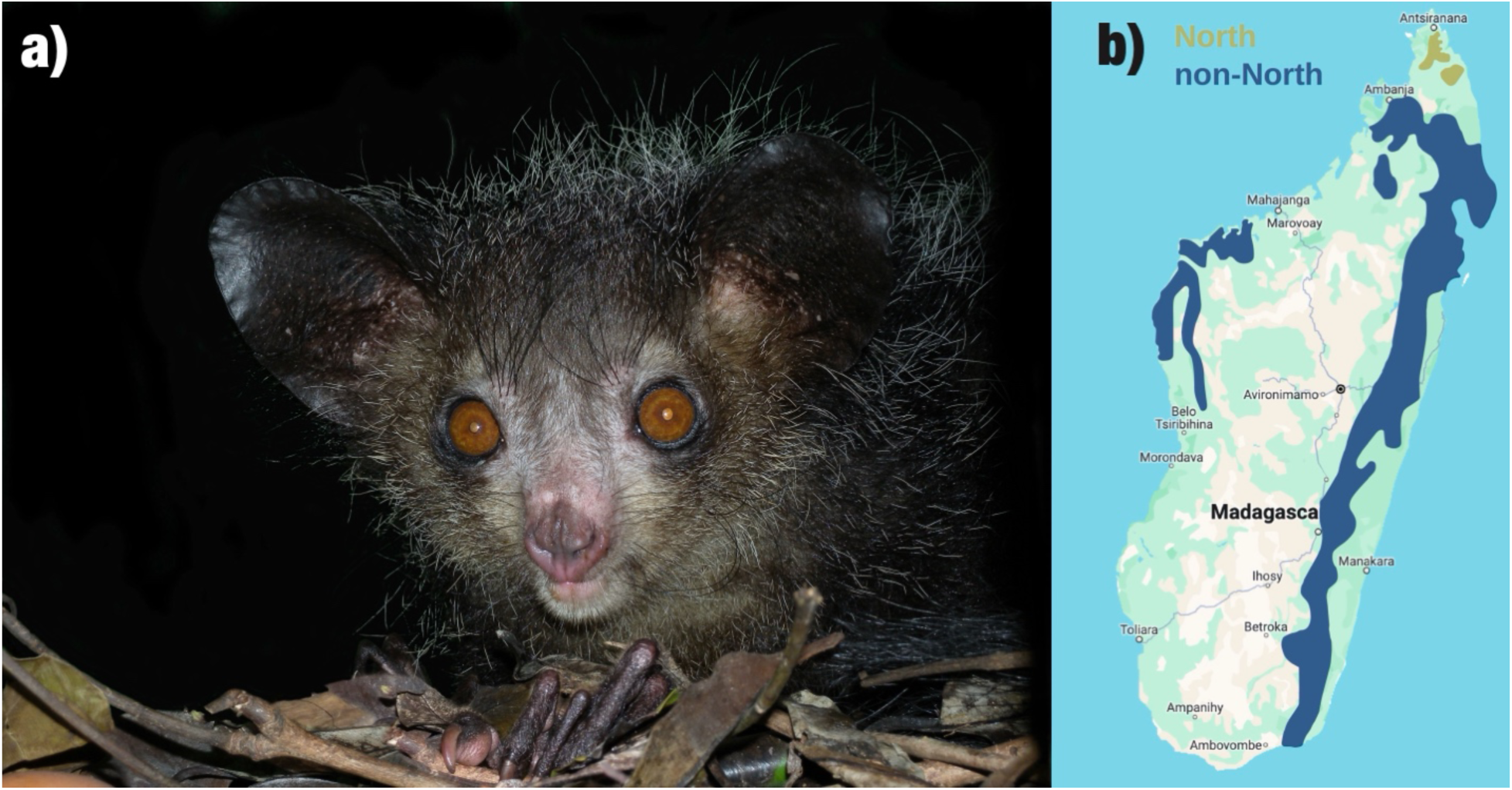
(a) An aye-aye from Madagascar; photo by Frank Vassen (Creative Commons CC BY 2.0 DEED). (b) Geographic distribution of the aye-aye (*Daubentonia madagascariensis*) in Madagascar, with populations in the Northern range shown in green and in the non-Northern range in blue. Range information was obtained from the International Union for the Conservation of Nature Annual Report 2020 (Louis et al. 2020).

The aye-aye’s range spans from the north to the south of the island of Madagascar (Figure 1b). Yet, as an arboreal primate, it is restricted to the western dry forests and the rainforests found in the east, central-west, and north of the island (Sterling 1994b; Louis *et al*. 2020). Additional populations were introduced to the nearby islands of Ile Roger and Nosy Mangabe off the coast of Madagascar in the 1960s (Petter 1977). However, one of the greatest threats to the survival of the species is persistent deforestation further reducing and fragmenting the limited viable ecosystems for the aye-aye (Suzzi-Simmons 2023). Such risk factors have contributed to the aye-aye and most other Malagasy lemur species being assessed as vulnerable or endangered by the International Union for the Conservation of Nature (IUCN; IUCN 2024).

Aye-aye population densities are generally thought to be very low (Mittermeier *et al*. 2010); however, as direct sightings of aye-ayes are rare, population studies on wild census populations typically rely on trace evidence such as feeding damage and tree holes (Sterling 1993; Ancrenaz *et al*. 1994; Sterling and Richard 1995; Randimbiharinirina *et al*. 2019). The IUCN has stressed an “urgent need for a systematic census of this important flagship species throughout its range” to allow for effective conservation efforts to be established (Louis *et al*. 2020). Provided that effective sampling and analyses are performed, population genomic assessments of the aye-aye’s historical range and population history can thus be essential for filling in these knowledge gaps.

The first population genomic studies performed over a decade ago found that aye-ayes have the lowest genetic diversity among extant lemur taxa, with estimates for the species as a whole ranging from 0.051% based on heterozygous sites in a single individual’s genome (Perry *et al*. 2012a) to 0.073% based on synonymous sites within transcriptome data of 1,175 genes from two individuals (Perry *et al*. 2012b). Genetic diversity within specific sub-populations corresponds to these estimated ranges with the Northern (*n* = 4), Eastern (*n* = 5), and Western (*n* = 3) sub-populations being 0.054%, 0.057%, and 0.049% respectively, based on synonymous sites in low-coverage sequencing data (Perry *et al*. 2013). These low estimates of genetic diversity further highlight the threat to the long-term prospects of this species.

Contemporary methods of demographic estimation empower us to further evaluate the status of the aye-aye, particularly when combined with high-quality, population-level, whole-genome sequencing data, as we present here. Equally importantly, the recent publication of a fully annotated, chromosome-level genome assembly of the aye-aye (Versoza and Pfeifer 2024) allows us to separate genomic data between functional / constrained sites and putatively neutral sites. This division is essential for inferring population history whilst avoiding mis-inference owing to Hill-Robertson effects (Hill and Robertson 1966); in particular, given that the vast majority of fitness-impacting mutations are deleterious, background selection (BGS) effects (Charlesworth *et al*. 1993) are expected to be relatively common across the genome (Charlesworth and Jensen 2021).

Two broad types of approaches have been proposed for dealing with the confounding effects of direct selection and selection at linked sites on demographic inference. In coding-dense genomes (as in many primate pathogens, for example), one would expect overlapping effects of both demography and selection across entire genomes (*e.g*., Irwin *et al*. 2016; Sackman *et al*. 2019; Jensen 2021; Morales-Arce *et al*. 2022; Terbot *et al*. 2023a,b; Howell *et al*. 2023; Soni *et al*. 2024a). In other words, because of the large genomic proportion of directly selected sites, there are likely no, or few, genomic regions that are free from selective effects. In such species, the joint and simultaneous inference of population history with selection becomes necessary, and approximate Bayesian approaches have been developed for this purpose (Johri *et al*. 2020, 2021, 2023). However, owing to the complexity of this joint inference framework, these approaches remain limited to highly simplified demographic models.

Conversely, in coding-sparse genomes (such as those characterizing aye-ayes and other primates), so-called “two-step” inference approaches are possible (see Soni and Jensen 2024b). Here, by first focusing on neutral sites at sufficient distances from functional sites in order to avoid these BGS effects, one may infer a neutral population history. Multiple well-performing inference approaches have been developed for such neutral demographic estimation (*e.g.*, Gutenkunst *et al*. 2009; Excoffier *et al*. 2013; and see the review of Beichman *et al*. 2018). In the second step, one may next study selective effects at and around functional sites, within the context of this inferred demography. Importantly, this framework for developing an appropriate evolutionary baseline model — a key component of which is the history of population size change, structure, and gene flow characterizing the species in question — has been repeatedly shown to be essential for reducing false-positive rates when scanning for loci targeted by recent positive or balancing selection (*e.g.*, Barton 1998; Jensen *et al*. 2005; Poh *et al*. 2014; Matuszewski *et al*. 2018; Harris and Jensen 2020; Charlesworth and Jensen 2022; Jensen 2023; Soni *et al*. 2023; Soni *et al*. 2024b; Soni and Jensen 2024a; and see the reviews of Johri *et al*. 2022a,b).

Improving the understanding of the aye-aye’s population dynamics will allow us to not only further assess the conservation status and evolutionary history of this unique species, but will also enable future study of the selective dynamics of the species as well. Additionally, the genomic resources resulting from these population-level sequence analyses will allow aye-ayes to serve as an invaluable outgroup for future primate analyses concerned with the study of virtually any evolutionary process or outcome, be it mutation, recombination, or distributions of fitness effects, to name but a few.

## RESULTS AND DISCUSSION

### Whole-genome, population-level data

To study the demographic history of the aye-aye, we combined publicly available low-coverage (7.1–10.6x) whole-genome sequencing data for 12 wild-caught individuals (Perry *et al*. 2013) with the genomes of five unrelated individuals housed at the Duke Lemur Center (DLC) that were newly sequenced at high-coverage (>50x) for this study (Supplementary Table 1). We mapped reads originating from each individual to the recently released high-quality, chromosome-level reference assembly for the aye-aye (Versoza and Pfeifer 2024), noting a much-improved mapping performance compared to the earlier reference assembly (Perry *et al*. 2012a) for the previously collected samples (94.5% compared to 84.9%; see Supplementary Table 1 in Perry *et al*. 2013). We subsequently called both variant and invariant sites according to the standard quality-control procedures outlined in the Genome Analysis Toolkit (GATK)’s Best Practices for non-model organisms (van der Auwera and O’Connor 2020). Following recommended guidelines in the field (Pfeifer 2017), we additionally excluded regions prone to misalignment and poor sequencing, yielding a dataset of 550,630 biallelic, autosomal single nucleotide polymorphisms (SNPs) with a transition-transversion ratio of 2.47 in the accessible genome (Supplementary Table 2). Lastly, as the effects of both direct and background selection have been shown to bias demographic inference (*e.g.*, Ewing and Jensen 2014, 2016; Johri *et al*. 2020, 2021; and see the reviews of Charlesworth and Jensen 2021, 2024), we limited this dataset to putatively neutral regions sufficiently distant from functional sites prior to any downstream analyses.

### Preliminary estimation of population structure and demography

Using marginal likelihood values produced by fastSTRUCTURE, population structure between two demes was found to be most likely (*k* = 1, *L* = −0.92860; *k* = 2, *L* = −0.90683; *k* = 3, *L* = −0.90684; *k* = 4, *L* = −0.90873; *k* = 5, *L* = −0.90685; with *k* being the number of demes and *L* the marginal likelihood). The assignment of individuals to their respective demes based on admixture proportions was almost complete (Figure 2), with all individuals having a 0.999998– 0.000002 deme assignment. Even for the second most likely structure of three demes, this level of assignment was maintained, with only two demes having individuals assigned to them by majority admixture proportion, and the third deme constituting a “ghost” population. The assignment of individuals used in Perry *et al*. (2013) for two demes matched those previously reported in constituting “North” and “non-North” demes, and all newly sequenced individuals were assigned to the non-North deme. While previous studies found support for a separation between Eastern and Western populations, this study did not find sufficient evidence to support this further division.

**Figure 2.**
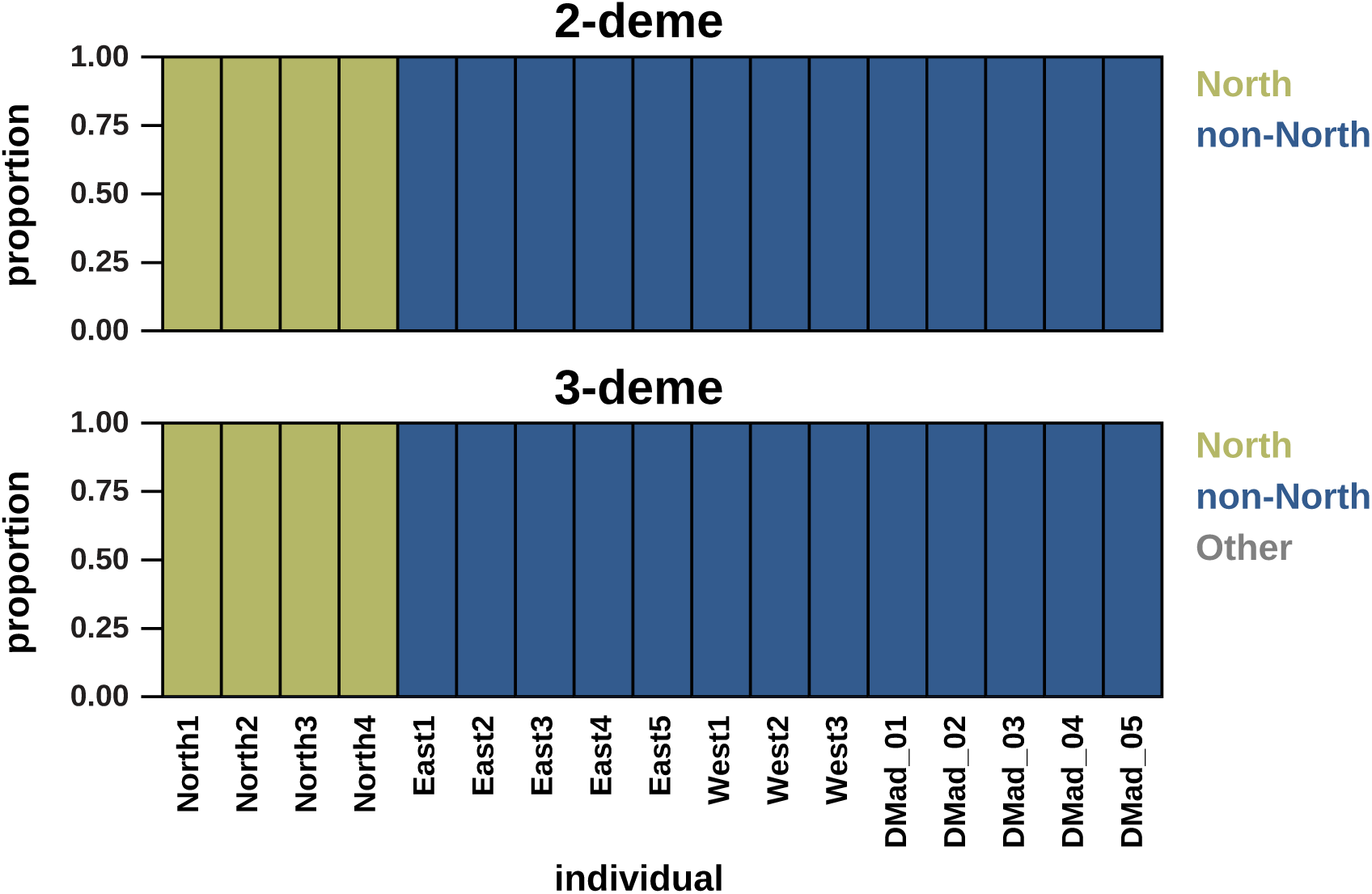
Admixture plots based on the results obtained from fastSTRUCTURE. Each individual is represented by a colored bar representing the demes (with the North population shown in green, the non-North population in blue, and the “ghost” population in gray).

Preliminary demographic history estimated using MSMC2 was characterized by a severe bottleneck deep in the aye-aye’s past, accompanied by a population split and recent rapid population growth, based upon the inclusion of either all invariant sites or only invariant sites within putatively neutral regions (Supplementary Figure 1). The estimated ancestral population size was approximately 340,000 when all invariant sites were used and 650,000 when only putatively neutral invariant sites were included. The estimated time of the bottleneck and population split occurred ∼8,370 generations ago, resulting in a North deme of ∼7,400 individuals and a non-North deme of ∼4,000 individuals, based upon putatively neutral invariant sites. The timing of this bottleneck when using all invariant sites was around 6,000 generations ago, and resulted in a North deme of ∼1,000 individuals and a non-North deme of ∼2,000 individuals. Under this inferred model, the population size of both demes would further reduce over time, and then experience recent rapid growth (within the past ∼3 generations) bringing population sizes back into the hundreds of thousands. This estimated rapid growth with large contemporary population sizes does not correspond well with our current understanding of aye-aye demographics, which is thought to be characterized by small and possibly declining population sizes (Louis *et al*. 2020). Importantly however, this U-shaped demographic pattern of ancient bottleneck and recent, explosive growth is a known mis-inference problem with MSMC2, and has been observed in a wide variety of organisms including vervet monkeys, passenger pigeons, elephants, *Arapidopsis*, and grapevines (see discussion of Johri *et al*. 2021, and additional performance issues resulting in erroneous population histories described in Mazet *et al*. 2016; Chikhi *et al*. 2017; Beichman *et al*. 2017). Notably, this MSMC2 estimated model indeed does not recapitulate our observed empirical data (see “Assessment of estimations using simulations” below).

### Demographic estimation using fastsimcoal2 and *δaδi*

Of the 16 models tested using fastsimcoal2, by far the most likely, based on maximum likelihood comparisons, was the bottleneck + decline model without migration (maximum likelihood = −3,406,939, Figure 3; Supplementary Table 3). The next most likely models were the bottleneck + growth without migration model (maximum likelihood = −3,407,054; Supplementary Figure 2) and the bottleneck without migration model (maximum likelihood = −3,407,079; Supplementary Figure 3). These results remained consistent when model comparisons were performed using the Akaike Information Criterion (AIC; Akaike 1974), which uses maximum likelihood scores alongside a penalty for the number of parameters in each model (ι1AIC for bottleneck + decline = 0.0, ι1AIC for bottleneck + growth = 229.1, and ι1AIC for bottleneck = 275.9). Despite seemingly large differences in the probabilities of each of these models, the estimated parameters for the bottleneck were surprisingly consistent: from an ancestral population size of ∼23,500 haploid genomes (*i.e.,* 11,750 diploid individuals), which were reduced to roughly 38% of that size ∼1,100 generations prior. More specifically, the best parameters for the bottleneck + decline model were an ancestral haploid effective population size of 23,389, experiencing a bottleneck 1,133 generations prior, reducing the total haploid size to 9,276, with 29% of the population in the North deme and the remainder in the non-North deme; more recently, these demes experienced further population decline at a constant rate resulting in an additional reduction in population sizes (to 525 diploid individuals in the North deme, and 1,285 diploid individuals in the non-North deme). Notably, based on a generation time of 5 years (Louis *et al*. 2020), this would place the initial bottleneck as occurring around 5,500 years before present, corresponding with the estimated arrival of humans to Madagascar (Dewar and Richard 2012; Salmona *et al*. 2017; Hansford *et al*. 2020; Balboa *et al*. 2024, but see Mitchell 2019). Moreover, the recent population decline in the past 35 years corresponds to observational reports of a ∼50% reduction in aye-aye populations during the past 20-40 years (Louis *et al*. 2020).

**Figure 3.**
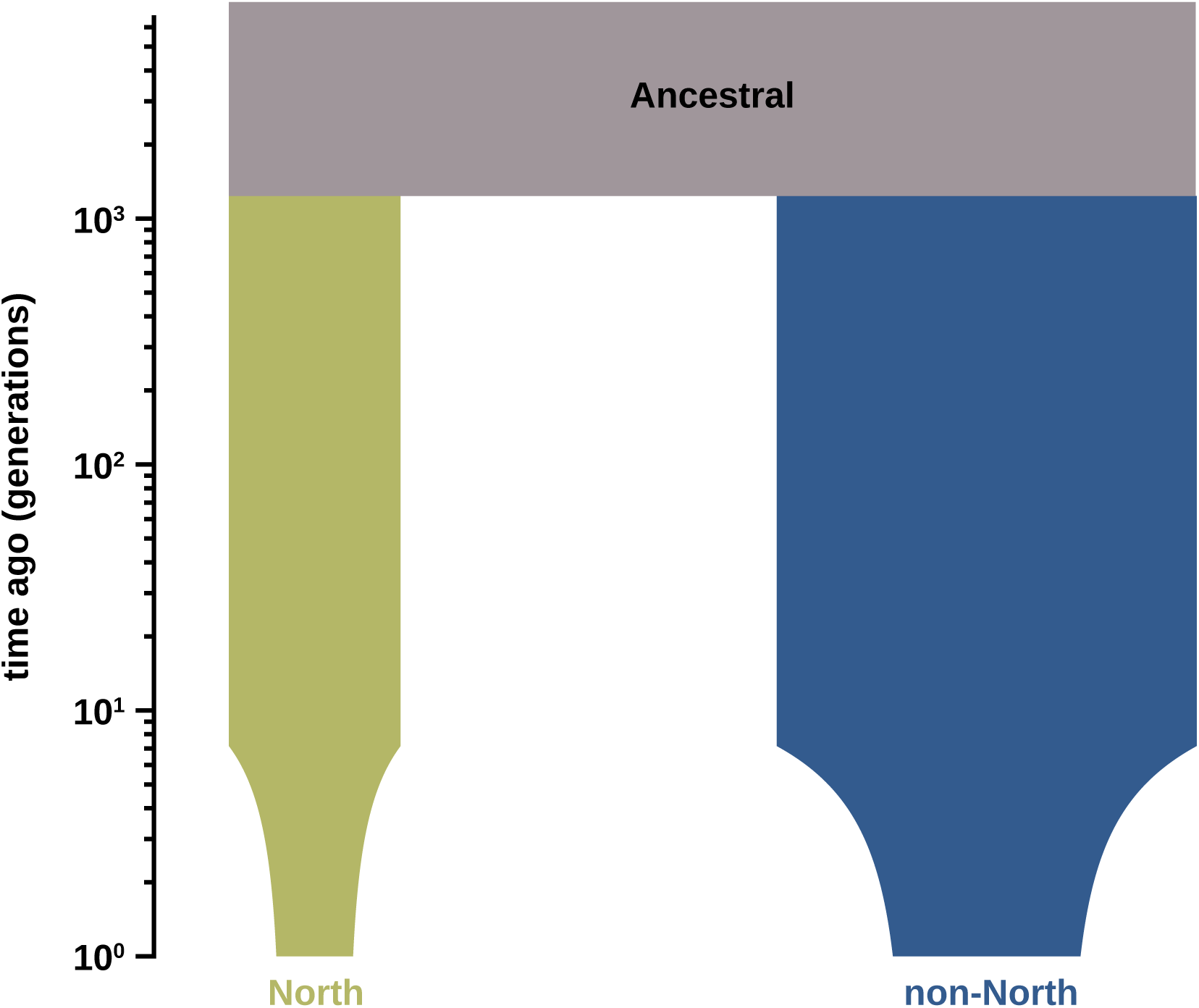
Diagram of the best-fitting demographic model produced by fastsimcoal2. The North population is shown in green, the non-North population in blue, and the ancestral population in taupe.

For comparative inference with *δaδi,* we ran the Portik *et al*. (2017) dadi_pipeline across 17 demographic models to identify the most appropriate (see the “Materials and Methods” section for more details). The *bottleneck_split_sizeChange* model had the highest mean log-likelihood (−1,420.676) and AIC (2,853.352) scores across five rounds of inference using dadi_pipeline. Notably, this model is similar to the best-fitting model described above as identified by fastsimcoal2. To infer the best-fitting parameters of this model, an additional 1,000 inference rounds were run. The inferred parameter values were broadly in agreement with the fastimcoal2 inference results, with an ancestral haploid population size of 23,000 (11,500 diploid individuals), and a bottleneck occurring 1,263 generations ago, resulting in daughter populations of size 3,152 haploid genomes (1,576 diploid individuals) in the North population and 8,174 haploid genomes (4,087 diploid individuals) in the non-North population. These populations then underwent a gradual population decline starting 40 generations ago, to estimated current day haploid genome sizes of 1,622 (811 diploid individuals) and 1,344 (672 diploid individuals) for the non-North and North populations, respectively. Supplementary Figures 4-5 provide a schematic of the best-fitting model and parameters, the comparative fit of the empirical and inferred two-dimensional side-frequency spectrum (SFS), and the residuals.

### Assessment of estimations using simulations

The fit of the SFS resulting from the estimated MSMC2 model (Supplementary Figure 6) to the observed data was extremely poor (Supplementary Figure 7), and this estimate can thus comfortably be disregarded as a viable model. By contrast, the resulting SFS for both the best fastsimcoal2 and *δaδi* models corresponded quite well to the observed SFS (Figure 4; Supplementary Figure 8). Furthermore, re-estimated parameters using the simulated SFS as input corresponded well to the distribution of estimated parameters using the observed SFS (Supplementary Figures 9 and 10), confirming model identifiability. For all estimated parameters, the ranges of those based on observed compared with those based on simulated data overlapped heavily. Together, this indicates that the estimated model and parameters are robust and consistent with observed genetic variation.

**Figure 4.**
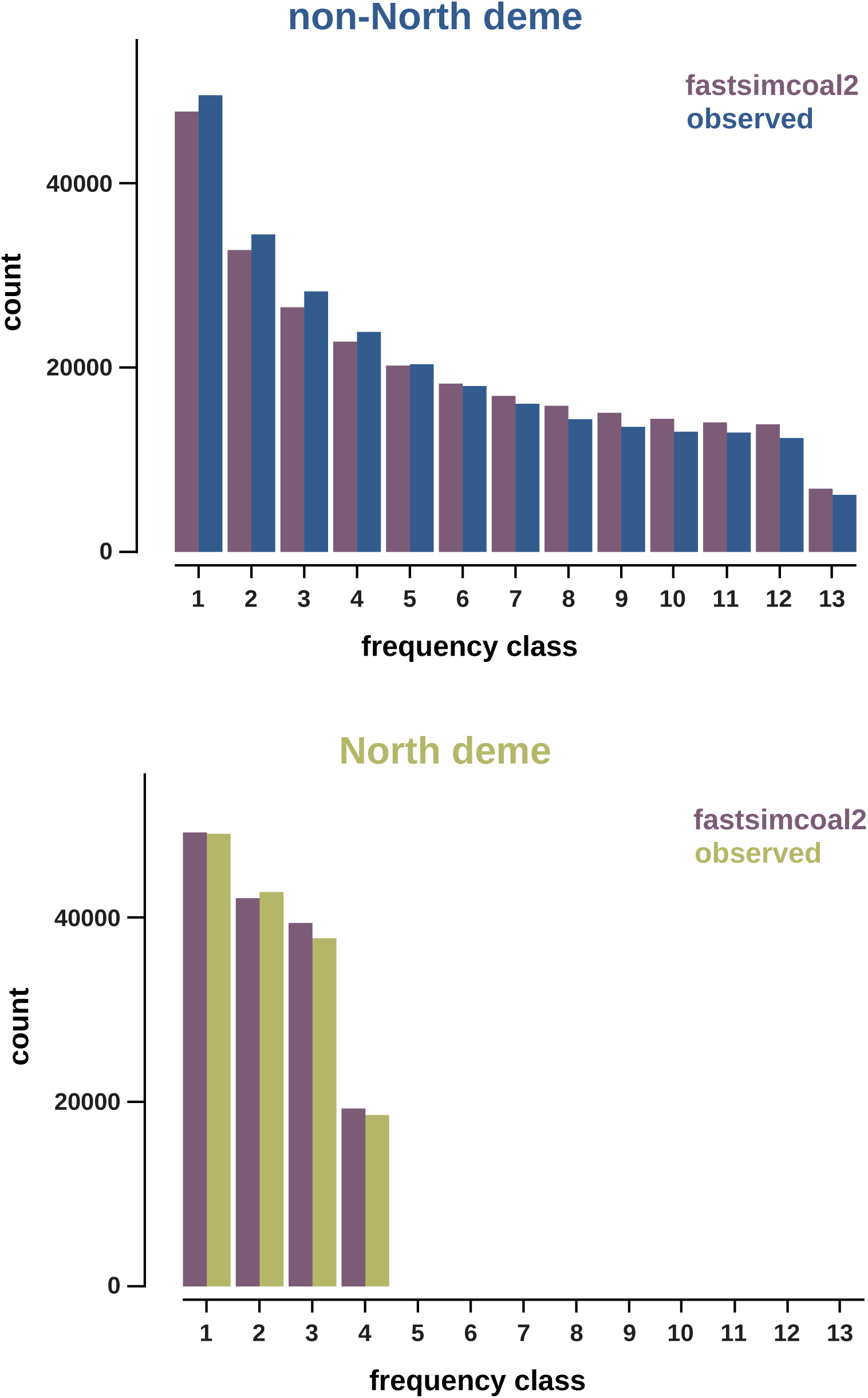
Folded site frequency spectra (SFS) of the best-fitting demographic models produced by fastsimcoal2, compared to the empirically observed SFS.

Overall, the similar results between fastsimcoal2 and *δaδi* are reassuring. Both approaches inferred models of population splits via bottleneck of approximately the same time period, followed by a recent population decline. This is particularly encouraging given that the *δaδi* model was uniquely able to include the possibility of exponential decline or growth. Both methods also inferred similar ancestral and post-bottleneck population sizes, as well as the timing of the population split.

## CONCLUSIONS

Inference suggests that the demographic history of the aye-aye is largely tied to the impacts of human activity. This study is consistent with field-based estimates suggesting a very recent decline of ∼50% in aye-aye populations occurring within the past 30-50 years. Additionally, we have found evidence for a prior bottleneck of ∼50% from the ancestral size that occurred 1,000-1,100 generations in the past. A population split into two demes (North and non-North) further accompanied this ancestral bottleneck, and is consistent with previously described population structuring (Perry *et al*. 2013). With an aye-aye generation time of 3-5 years, this corresponds well with the estimated timing of the first human settlements on the island of Madagascar, though that colonization date is still a topic of considerable uncertainty with estimates ranging from 1,000 to 10,000 years before present (Hansford *et al*. 2020). Notably, similar population declines and fragmentations for at least two other species of lemur on Madagascar have been dated to the same general period: the golden crowned sifaka (*Propithecus tattersalli*) and Perrier’s sifaka (*P. perrieri*) (Salmona *et al*. 2017). In addition, demographic estimates of Malagasy populations of the African bush pig (*Potamochoerus larvatus*) include a founding bottleneck also dating to the same period, indicating a possible introduction to the island by humans (Balboa *et al*. 2024). While climatic changes not driven by human land use may have contributed to the population fragmentation and decline of lemur species on Madagascar over the past several thousand years (Salmona *et al*. 2017), the impact of humans on Malagasy wildlife, particularly in the past century, is increasingly well supported by such estimates of demographic history. Given that human arrival is generally accepted to impact the population sizes of local megafauna, the incidental evidence from these demographic studies may even be interpreted as providing support for dating the arrival of humans to Madagascar to 3,000 to 5,000 years before present.

Regarding future work on the aye-aye, this study supports two main points. First, conservation measures are at a critical point as the population size of the aye-aye appears to be in decline, with the total effective population size now estimated to be below 2,000 individuals. Second, the well-fitting demographic history inferred here will prove valuable in all future population genomic studies in the species, as this neutral baseline model — accounting for the dominant role of population history in shaping allele frequencies — is a necessary pre-requisite for further inference ranging from the detection of selective sweeps to population-level estimates of genome-wide recombination rates (Johri et al. 2022a).

## MATERIALS AND METHODS

### Animal subjects

The DLC’s Research Committee (protocol BS-3-22-6) and Duke University’s Institutional Animal Care and Use Committee (protocol A216-20-11) approved this study. We performed this study in compliance with all regulations regarding the care and use of captive primates, including the U.S. National Research Council’s Guide for the Care and Use of Laboratory Animals and the U.S. Public Health Service’s Policy on Human Care and Use of Laboratory Animals.

### Sample collection, preparation, and sequencing

We extracted genomic DNA (gDNA) from whole blood samples obtained from five aye-ayes (*D. madagascariensis*) housed at the DLC (Durham, NC, USA) – two wild-born females, two wild-born males, and one colony-born female of wild-born parents – using the PureLink Genomic DNA Mini Kit. For each sample, we prepared a 150 bp paired-end library using the NEBNext Ultra II DNA PCR-free Library Prep Kit (New England Biolabs, Ipswich, MA, USA). We checked qualities and quantities of libraries using a High Sensitivity D1000 ScreenTape on an Agilent TapeStation (Agilent Technologies, Palo Alto, CA, USA), a Qubit 2.0 Fluorometer (ThermoFisher Scientific, Waltham, MA, USA), and real-time PCR (Applied Biosystems, Carlsbad, CA, USA) prior to sequencing them on the Illumina NovaSeq platform (Illumina, San Diego, CA, USA) to >50x coverage each (Supplementary Table 1).

Additionally, we downloaded publicly available, low-coverage (7.1–10.6x) whole-genome sequencing data for 12 wild-caught individuals from the NCBI Sequence Read Archive (BioProject PRJNA189299). This data was previously generated by extracting gDNA from samples of liver tissue and whole blood using the Illumina TruSeq DNA Sample Preparation v2 Low Throughput Protocol, shearing the DNA to ∼300 bp fragments using a Covaris S2 instrument, enriching the fragments via PCR, and 101 bp paired-end sequencing on the Illumina HiSeq 2000 platform (Illumina, San Diego, CA, USA) (for details, see Perry *et al*. 2013).

### Variant calling, genotyping, and filtering

To avoid technical artifacts, we first trimmed raw reads using TrimGalore v.0.6.10 (https://github.com/FelixKrueger/TrimGalore) with a default quality trimming threshold of 20, and we marked the locations of adapter sequences in both the novel and public whole-genome data using GATK *MarkIlluminaAdapters* v.4.1.8.1 and v.4.2.6.1 (van der Auwera and O’Connor 2020), respectively. We then used the Burrows Wheeler Aligner (BWA-MEM) v.0.7.17 (Li and Durbin 2009) to map the pre-processed reads to the aye-aye reference assembly (DMad_hybrid; GenBank accession number: JBFSEQ000000000; Versoza and Pfeifer 2024) and identified duplicate reads using GATK’s *MarkDuplicates* v.4.1.8.1 and v.4.2.6.1. We refined read alignments by performing multiple sequence realignments around insertions and deletions using GATK’s *RealignerTargetCreator* and *IndelRealigner* v.3.8, recalibrating base quality scores using GATK’s *BaseRecalibrator* and *ApplyBQSR* v.4.2.6.1 together with high-confidence training data obtained from pedigreed individuals housed at the DLC (Versoza *et al*. 2024a,b), and, finally, carrying out a further round of duplicate removal using GATK’s *MarkDuplicates* v.4.2.6.1. We then used high-quality read mappings (‘*--minimum-mapping-quality* 40’) to call variant and invariant sites using GATK’s *HaplotypeCaller* v.4.2.6.1 in the ‘*-ERC* BP_RESOLUTION’ mode, with the ‘*--heterozygosity*’ parameter set to 0.0005 to reflect the level of genetic diversity previously observed in the species (Perry *et al*. 2013). To account for differences in the library preparation protocols (see “Sample collection, preparation, and sequencing”), we set the ‘*--pcr-indel-model*’ to NONE for the novel data (five individuals) and CONSERVATIVE for the previously published data (12 individuals). We merged individual calls using GATK’s *CombineGVCFs* v.4.2.6.1 and jointly genotyped them at all sites (‘*-all-sites*’) using GATK’s *GenotypeGVCFs* v.4.2.6.1.

We partitioned the call set into autosomal, biallelic SNPs and invariant sites genotyped in all individuals (AN = 34) and filtered sites according to the GATK Best Practices “hard filter” criteria (*i.e.*, QD < 2.0, QUAL < 30.0, SOR > 3.0, FS > 60.0, MQ < 40.0, MQRankSum < −12.5, and ReadPosRankSum < −8.0; with acronyms as defined by the GATK package) using GATK’s *SelectVariants* and *VariantFiltration* v.4.2.6.1, respectively. Additionally, we excluded regions prone to misalignment and/or poor sequencing by removing repetitive elements annotated in the DMad_hybrid genome assembly (Versoza and Pfeifer 2024) and applying upper and lower cutoffs on the individual depth of coverage (0.5 × DP_ind_ and 2 × DP_ind_).

### Putatively neutral regions

Prior to performing demographic inference, we limited the dataset to putatively neutral regions in order to avoid the confounding effects of direct selection, or selection at linked sites. To this end, following Johri *et al*.’s (2020, 2023) recommendations for constructing a species-specific evolutionarily appropriate null model, we used the DMad_hybrid genome annotation (Versoza and Pfeifer 2024), containing a total of 33,346 genes (18,858 of which protein-coding), to remove any sites falling within 10 kb of exons. This trimming was intended to largely eliminate the effects of background selection, which has been shown to bias demographic inference (*e.g.*, Ewing and Jensen 2014, 2016; Johri *et al*. 2020, 2021; and see the reviews of Charlesworth and Jensen 2021, 2024). Moreover, as additional sites may be experiencing purifying selection (*e.g.*, RNA binding sites; Simkin *et al*. 2014), sequence elements previously identified as constrained across 239 primate genomes (Kuderna *et al*. 2024) were excluded from further analyses. For simplicity, we will refer to the excluded sites as “selectively constrained” and the included sites as “putatively neutral” elsewhere in this study.

### Preliminary estimation of population structure and demography

As our initial step in estimating the demographic history of the aye-aye, we determined the extent of population structure within our sample. To this end, we first limited our dataset to putatively neutral SNPs using VCFtools v.0.1.16 (Danecek *et al*. 2011) and converted the output to the format required by fastSTRUCTURE (Raj *et al*. 2014) using PGDSpider v.2.1.1.5 (Lischer and Excoffier 2012). Using this dataset, we assessed population structure using the independent-loci model implemented in fastSTRUCTURE v.1.0, thereby allowing individuals to exhibit ancestry from one to five demes (*i.e.*, deme counts ranged from *k* = 1 to *k* = 5). From these results, we then selected the number of demes and deme assignment of individuals based on the value of *k* (deme count) that had the greatest maximum likelihood.

Following this, we processed the sequence information along with filters detailing selectively constrained regions and total site counts which passed quality filtering using a custom Perl script to generate input files for MSMC2 (Schiffels and Durbin 2015) which contained information on putatively neutral SNPs and the number of intervening invariant sites. Invariant sites within selectively constrained regions were alternatively included or silenced from analyses for comparison; in addition, only variant sites within the putatively neutral regions that were either phased for all individuals or homozygous within all individuals were included. Using the deme assignment recommended by fastSTRUCTURE, we analyzed these input files with MSMC2 v.2.1.4 to provide an initial estimation of demographic history.

### Demographic estimation using fastsimcoal2

Based on the information obtained from the preliminary analyses using fastSTRUCTURE and MSMC2, we wrote several demographic models as input files for fastsimcoal2 (Excoffier *et al*. 2013, 2021; Marchi *et al*. 2023). Specifically, we tested four model sets, all with an asymmetric population split occurring in the past. One model set included a constant rate of population decline since the population split; others allowed a population bottleneck to occur alongside the population split followed by either no subsequent change in population size, a constant rate of recent population growth, or a constant rate of recent population decline. For all model sets, both demes shared the bottleneck ratio, growth rates, and decline rates when relevant. All sets included models with and without migration since the population split; we tested migration between the two demes at both asymmetric and symmetric rates. Note that in the context of fastsimcoal2 “population size” refers to the effective population size of haploid genomes and not necessarily precise census counts of diploid individuals.

The parameter search method used by fastsimcoal2 uses an initial range with a fixed lower bound as a minimum value and a flexible upper bound that can be extended unless explicitly told to use the provided upper bound as a fixed maximum value. Across the model sets tested, nine different parameters were estimated: current population size of the North deme (1,000 to 100,000, all models), current population size of the non-North deme (1,000 to 100,000, all models), time since recent recovery growth or population decline began (1 to 15 generations, bottleneck + growth and bottleneck + decline models), ratio between current and pre-growth or pre-decline population sizes (0.01 to 1 bounded, bottleneck + growth models; 1 to 3 bounded, bottleneck + decline models), time since bottleneck from last event (5 to 250 generations, bottleneck models), ratio between post- and pre-bottleneck population sizes (0.0001 to 0.5 bounded, bottleneck models), time since population decline began (1 to 250 generations, continuous decline models), ratio between pre-decline and current population sizes (1.1 to 100,000, continuous decline models), and migration rates (0.0001 to 0.1, migration models; two parameters with same initial ranges were used for models with asymmetric migration).

Once we prepared model input files, we produced the multidimensional SFS for the putatively neutral SNPs using easySFS v. 0.0.1 (Coffman *et al*. 2015) and the sequence data prepared previously. From this, we estimated model parameters using fastsimcoal2 v.28 with the following settings: 150,000 simulations per tested parameter, 50 maximization cycles, and resetting parameter searches after 10 consecutive failed cycles (-n 150000 -L 50 -y 10). We estimated parameters with 250 replicates for each model. We then selected the best performing model and parameter combinations based on maximum likelihood and minimizing the difference between estimated and observed likelihoods.

### Demographic estimation using *δaδi*

To further validate our demographic estimations, we additionally performed parameter estimation using *δaδi* (Gutenkunst *et al*. 2009). Specifically, we fit 17 two-population demographic models using dadi_pipeline v.3.1.5 (Portik *et al*. 2017). These models included simple models, simple models plus instantaneous size change, ancient migration or secondary contact, ancient migration or secondary contact plus instantaneous size change, and two-epoch models with continuous migration, as described in the dadi_pipeline documentation. Additionally, we added a model of our own, based on the best-fitting fastsimcoal2 model. This model – named *bottleneck_split_sizeChange* – consists of a two-population split from the ancestral population via a bottleneck, followed by exponential size change in each population. Where the model differs from the best-fitting fastsimcoal2 model is that each daughter population was given an individual size change parameter, allowing for either exponential growth or decline. This model consisted of six parameters: *nuB1* (ratio of the population size of the non-North population relative to the ancestral population after the population split), *nuB2* (ratio of the population size of the North population relative to the ancestral population after the population split), *Ts* (time at which the population split occurred, measured in units of 2*N_a_* generations, where *N_a_* is the ancestral population size), *Tg* (time at which exponential growth/decline began in both daughter populations, measured in units of 2*N_a_* generations), *nuF1* (ratio of the population size of the non-North population relative to the ancestral population at the time of sampling)*, nuF2* (ratio of the population size of the North population relative to the ancestral population at the time of sampling). Following the dadi_pipeline guidelines, the best-fitting model was inferred based on the AIC score (Akaike 1974).

Once the best-fitting model was inferred (*bottleneck_split_sizeChange* - based on consistently lowest AIC values across multiple runs), we reran parameter inference on this model across 1,000 inference replicates, with the optimization function set to 300 maximum iterations, using starting parameter values of *nuB1* = 0.1; *nuB2* = 0.1; *Ts* = 0.05; *Tg* = 0.005; *nuF1* = 0.1; *nuF2* = 0.1, and using *δaδi*’s *perturb_params* function to adjust these parameters two-fold up or down, within the following lower and upper parameter bounds: 0.01 ≤ *nuB1* ≤ 0.5; 0.01 ≤ *nuB2* ≤ 0.5; 0.01 ≤ *Ts* ≤ 0.1; 0.001 ≤ *Tg* ≤ 0.01; 0.01 ≤ *nuF1* ≤ 0.5; 0.01 ≤ *nuF2* ≤ 0.5. The best-fit parameter combination was that which yielded the highest log-likelihood score.

### Assessment of inference using simulations

To assess the fit of our model selection and parameter estimation, we simulated 100 replicates of the best-fitting models and their parameters and visually compared the simulated to the observed SFS. For the best model-parameter combinations from fastsimcoal2 and MSMC2, we simulated SFS with fastsimcoal2; for the best model-parameter combination from *δaδi*, we performed simulations using msprime v. 1.3.2 (Baumdicker *et al*. 2022). Recombination rates were simulated to match previously observed pedigree data (1e-8 cM/Mb, Versoza, Lloret-Villas et al. 2024), and mutation rates were set to that previously reported in another lemur species (1.52e-8 per base pair per generation, Campbell *et al*. 2021; and as modeled in Soni *et al*. (2024c). For each of the simulated SFS, we estimated parameters using fastsimcoal2 and *δaδi* in a manner similar to the empirical data but performing only 100 (rather than 250) replicates per simulated SFS in fastsimcoal2. We then visually compared the distribution of estimated parameters determined using the observed SFS to the distribution of best estimated parameters based on the simulated SFS.

## ACKNOWLEDGEMENTS

We would like to thank Erin Ehmke, Kay Welser, and the Duke Lemur Center for providing the aye-aye samples used in this study, and members of the Jensen Lab and Pfeifer Lab for helpful comments and discussion. DNA extraction, library preparation, and Illumina sequencing was conducted at Azenta Life Sciences (South Plainfield, NJ, USA). Parameter estimations using fastsimcoal2 were performed using resources from the Open Science Pool (OSG 2006; Pordes *et al*. 2007; Sfiligoi *et al*. 2009; OSG 2015). Computations were performed on the Sol supercomputer at Arizona State University (Jennewein et al. 2023). This is Duke Lemur Center publication # XXXX.

