## Supplementary Materials for "Inferring the demographic history of aye-ayes (*Daubentonia madagascariensis*) from high-quality, whole-genome, population-level data"

### SUPPLEMENTARY TABLES AND FIGURES

| | NCBI | coverage |
|---|---|---|
| novel | DMad_01 | 104.9 |
| | DMad_02 | 50.5 |
| | DMad_03 | 50.2 |
| | DMad_04 | 53.7 |
| | DMad_05 | 52.5 |
| public | East1 | 10.6 |
| | East2 | 9.2 |
| | East3 | 10.2 |
| | East4 | 10.6 |
| | East5 | 9.0 |
| | North1 | 7.0 |
| | North2 | 9.5 |
| | North3 | 8.3 |
| | North4 | 7.4 |
| | West1 | 8.4 |
| | West2 | 7.1 |
| | West3 | 8.0 |

**Supplementary Table 1**. Samples and their sequencing coverage.

| scaffold | length | # SNPs | # invariant sites | Ts/Tv |
|---|---|---|---|---|
| 1 | 316,165,773 | 82,219 | 35,139,180 | 2.37 |
| 2 | 290,592,686 | 72,023 | 32,727,516 | 2.43 |
| 3 | 261,424,170 | 73,340 | 30,723,768 | 2.37 |
| 4 | 219,686,500 | 59,431 | 25,957,079 | 2.37 |
| 5 | 215,448,047 | 54,199 | 24,631,731 | 2.43 |
| 6 | 204,016,426 | 50,429 | 23,226,167 | 2.48 |
| 7 | 199,604,927 | 42,325 | 19,504,415 | 2.43 |
| 8 | 162,769,830 | 40,950 | 17,706,139 | 2.40 |
| 10 | 114,896,738 | 25,555 | 11,817,922 | 2.43 |
| 11 | 102,076,017 | 14,070 | 6,936,316 | 2.58 |
| 12 | 67,301,774 | 9,779 | 4,242,984 | 2.57 |
| 13 | 62,733,483 | 9,219 | 4,167,735 | 2.51 |
| 14 | 34,254,822 | 8,930 | 3,562,322 | 2.65 |
| 15 | 28,257,198 | 8,161 | 3,383,534 | 2.60 |
| ∑ or ∅ | **2,279,228,391** | **550,630** | **243,726,808** | **2.47** |

**Supplementary Table 2**. Summary of genomic data. After filtering, a total of 550,630 autosomal, biallelic, single nucleotide polymorphisms (SNPs) with a transition-transversion ratio (Ts/Tv) of 2.47 were discovered in the accessible genome.

| model name | *k* | max. likelihood | AIC | ∆AIC |
|---|---|---|---|---|
| 2-decline | 3 | -3407626 | 6815258 | 1368.939 |
| 2-decline-mig1 | 4 | -3412914 | 6825836 | 11947.24 |
| 2-decline-mig2 | 5 | -3408936 | 6817883 | 3993.608 |
| 2-bn | 3 | -3407079 | 6814165 | 275.8654 |
| 2-bn-mig1 | 4 | -3409659 | 6819327 | 5437.563 |
| 2-bn-mig2 | 5 | -3409827 | 6819665 | 5775.846 |
| 2-bn-recover | 5 | -3407054 | 6814118 | 229.0796 |
| 2-bn-recover-mig1 | 6 | -3409764 | 6819540 | 5651.195 |
| 2-bn-recover-mig2 | 7 | -3409084 | 6818182 | 4292.606 |
| 2-bn-recover-mig3 | 6 | -3409796 | 6819605 | 5715.593 |
| 2-bn-recover-mig4 | 7 | -3409404 | 6818822 | 4933.153 |
| 2-bn-decline | 5 | -3406940 | 6813889 | 0 |
| 2-bn-decline-mig1 | 6 | -3408673 | 6817358 | 3469.048 |
| 2-bn-decline-mig2 | 7 | -3408833 | 6817680 | 3791.384 |
| 2-bn-decline-mig3 | 6 | -3408760 | 6817531 | 3642.281 |
| 2-bn-decline-mig4 | 7 | -3409132 | 6818277 | 4388.338 |

**Supplementary Table 3**. Number of parameters (*k*), maximum likelihood, Akaike Information Criterion (AIC) score, and ∆AIC scores for the best parameters for each model tested using fastsimcoal2. The best fitting model tested based on both maximum likelihood and AIC was the model with a bottleneck + split ("bn"), recent decline ("decline"), and no migration (highlighted in green). Models with "mig1" or "mig3" modeled symmetrical migration rates, while those with "mig2" or "mig4" modeled asymmetric migration rates. Migration in "mig1" and "mig2" models occurred continuously since the population split, while migration in "mig3" and "mig4" models occurred only during recent population decline or growth ("recover"). The "2-decline" models did not include a bottleneck alongside the population split and instead modeled a constant rate of population decline since the split.

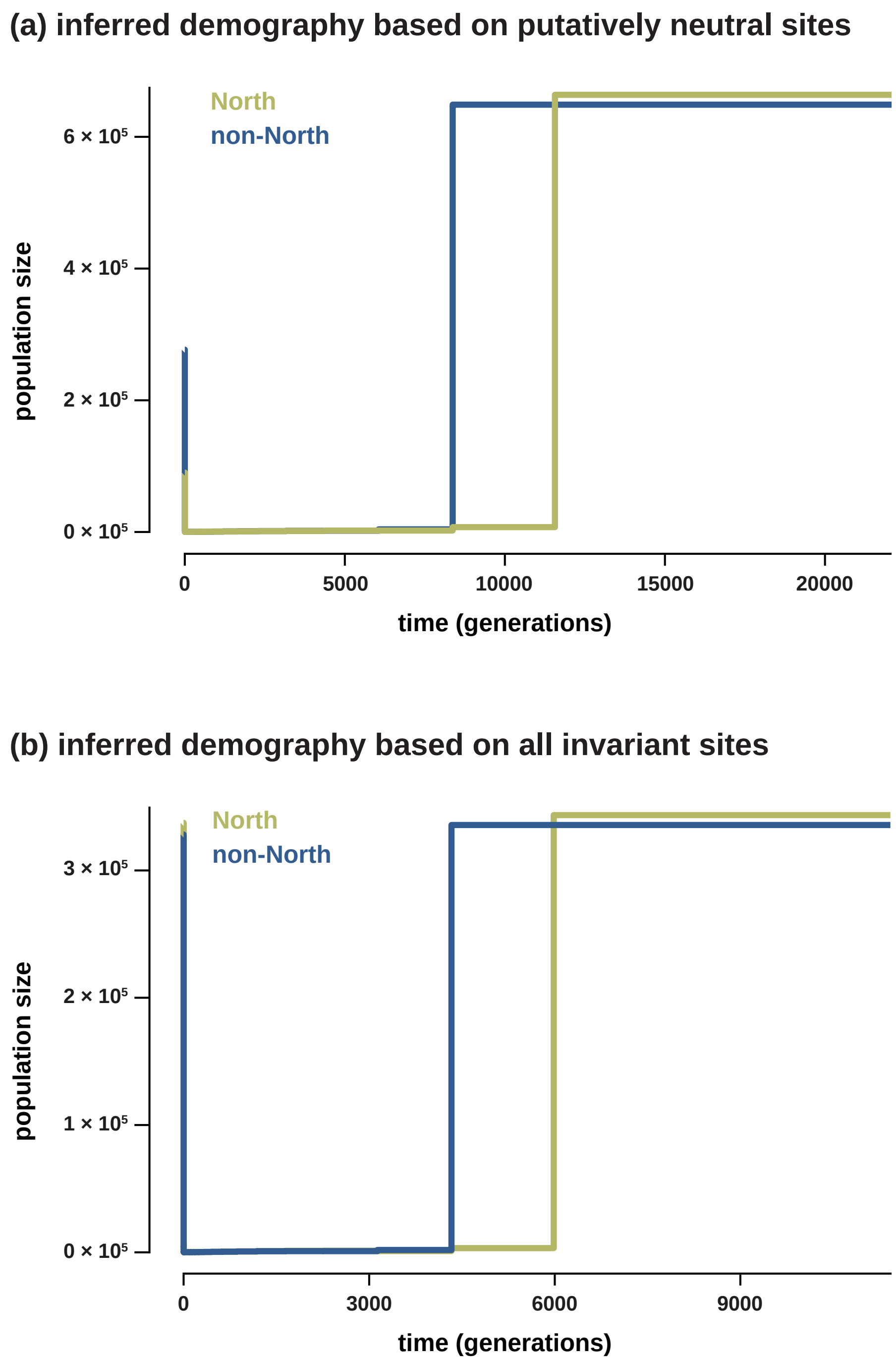


**Supplementary Figure 1**. MSMC2 plots of population size of the two demes (North and non-North shown in green and blue, respectively) over time using (a) variant and invariant sites from putatively neutral regions only, and (b) based on all invariant sites.

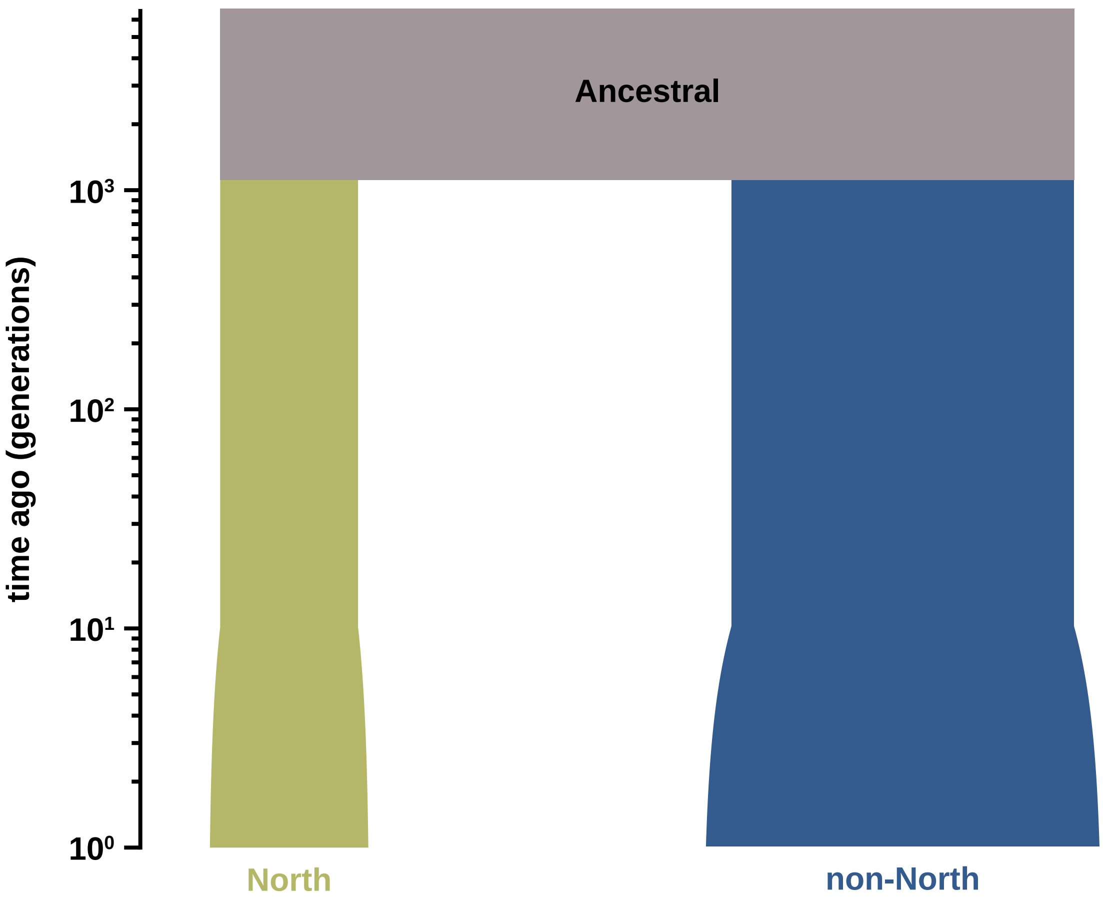


**Supplementary Figure 2**. Diagram of the best-fitting parameters produced by fastsimcoal2 for the bottleneck + split, recent growth, and no migration model. The North population is shown in green, the non-North population in blue, and the ancestral population in taupe.

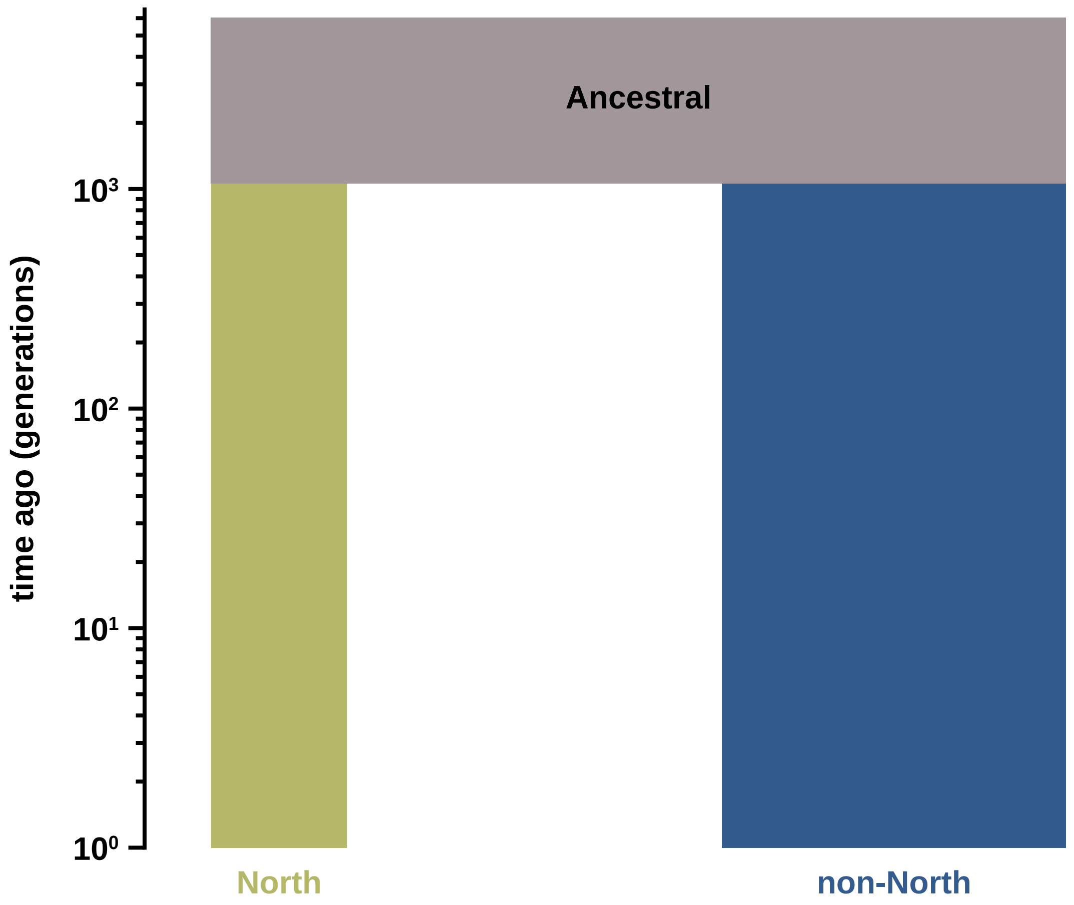


**Supplementary Figure 3**. Diagram of the best-fitting parameters produced by fastsimcoal2 for the bottleneck + split and no migration model. The North population is shown in green, the non-North population in blue, and the ancestral population in taupe.

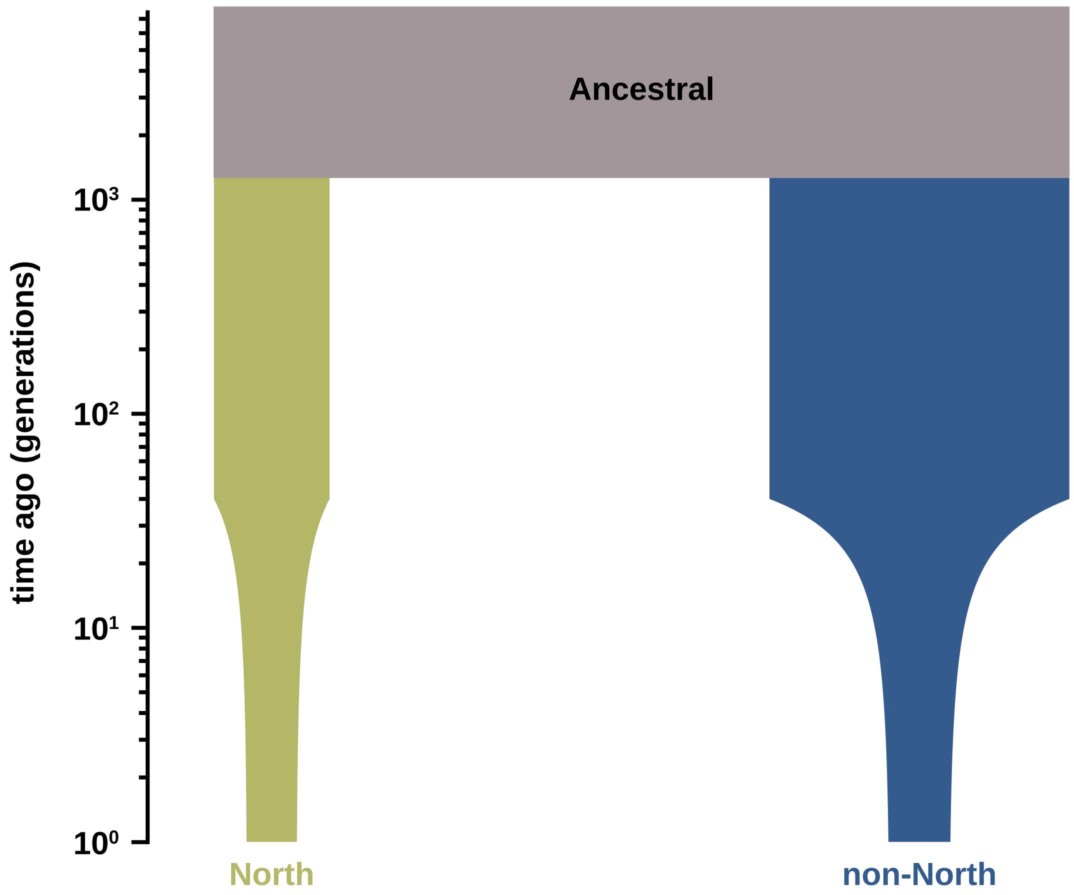


**Supplementary Figure 4**. Diagram of the best-fitting model and parameters produced by *δaδi*. The North population is shown in green, the non-North population in blue, and the ancestral population in taupe.

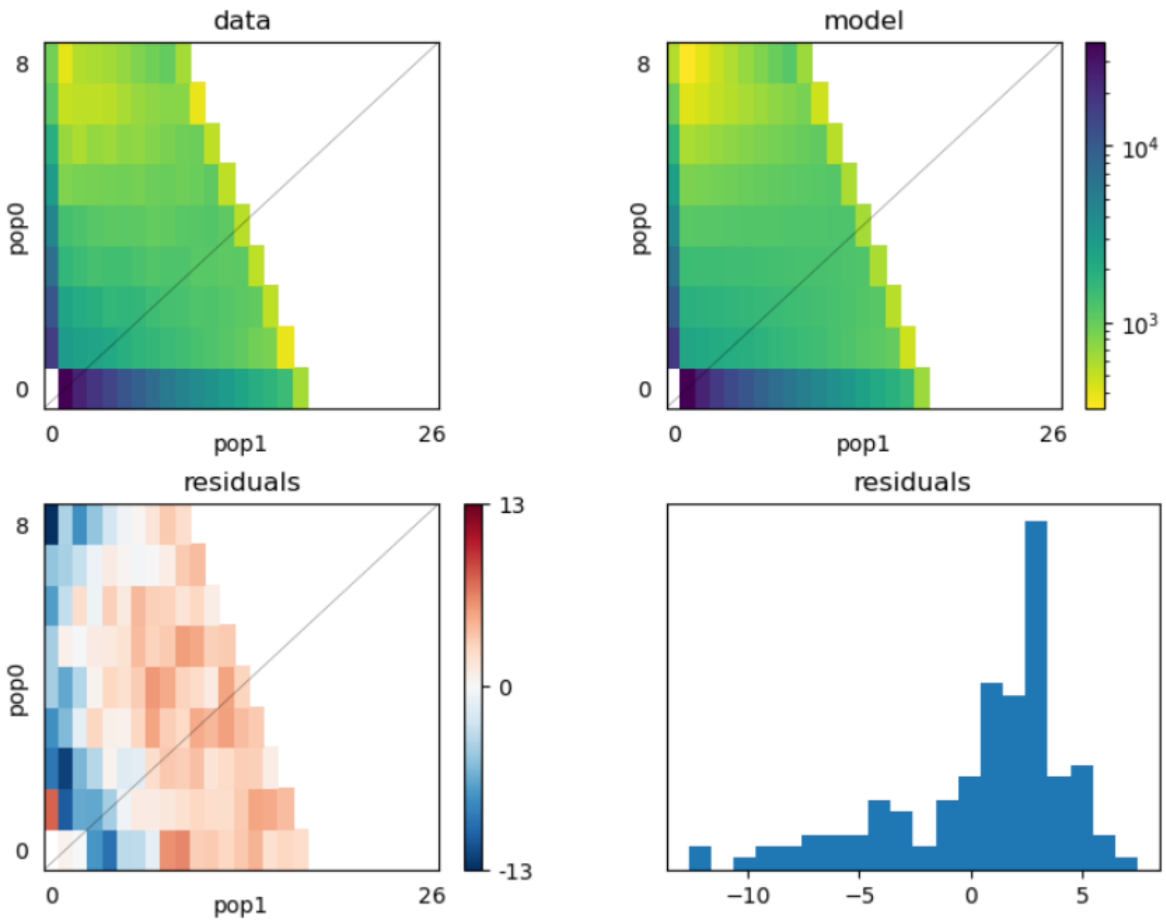


**Supplementary Figure 5**. Residuals corresponding to the best-fitting *δaδi* model and parameters, as presented in Supplementary Figure 4.

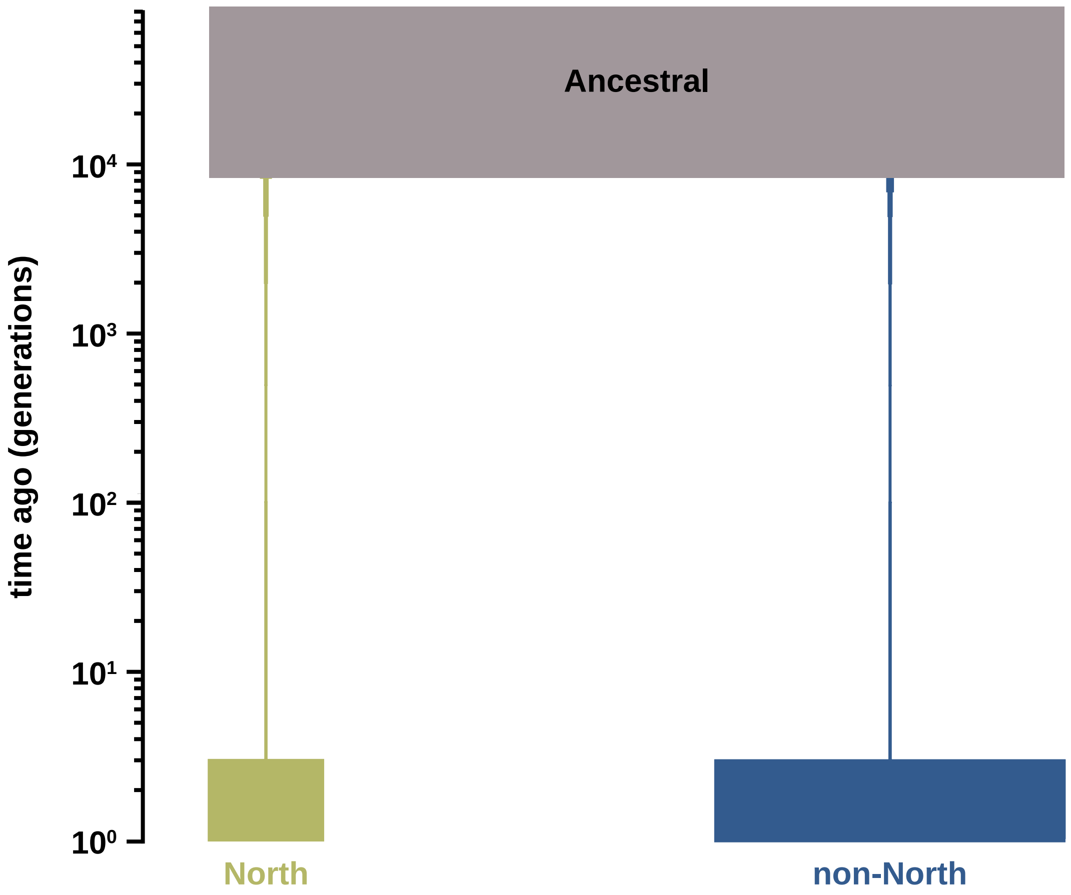


**Supplementary Figure 6**. Diagram of the approximation of the demographic history estimated by MSMC2. The North population is shown in green, the non-North population in blue, and the ancestral population in taupe.

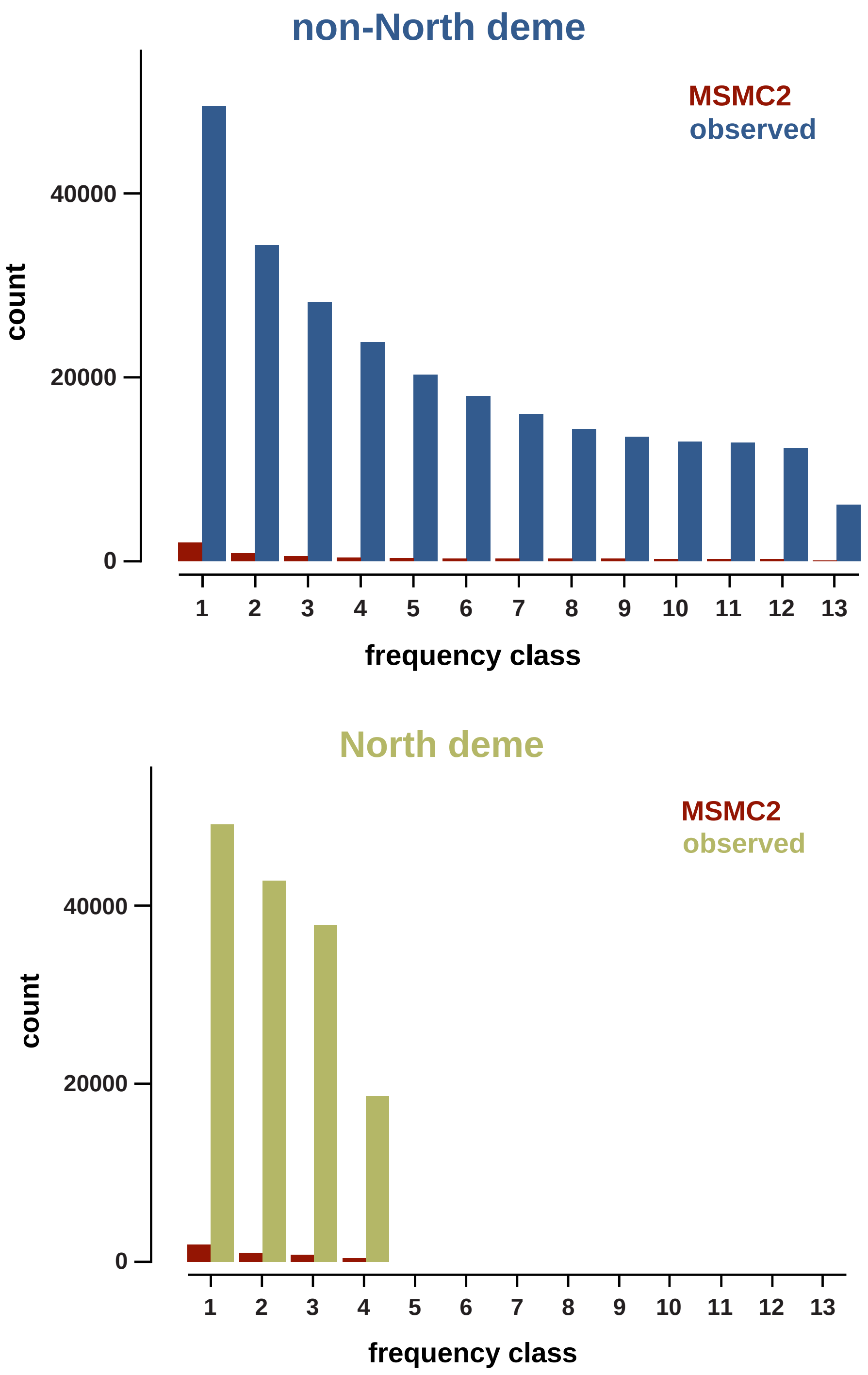


**Supplementary Figure 7**. Folded site frequency spectrum (SFS) of the best-fitting MSMC2 model vs. the observed SFS. Notably, the large number of 'missing' single nucleotide polymorphisms (SNPs) in the MSMC2 model relative to the observed data owes to the fact that, under this model, >2 million species-level SNPs become fixed differences between the North and non-North populations, and thus do not appear in the SFS as segregating variants in either population.

a)

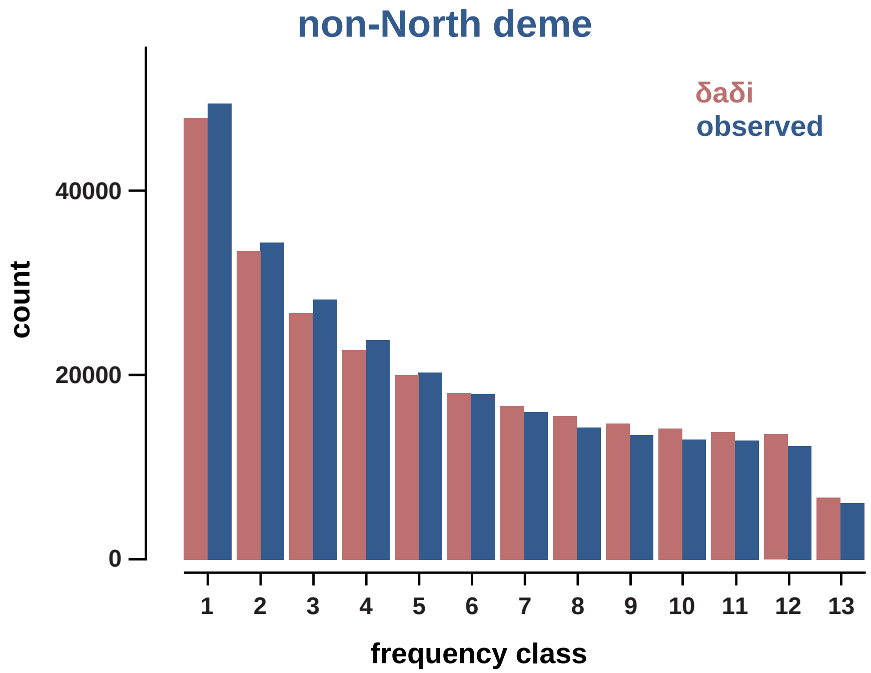

non-North deme
δaδi
observed
40000
20000
0
count
1
2
3
4
5
6
7
8
9
10
11
12
13
frequency class


b)

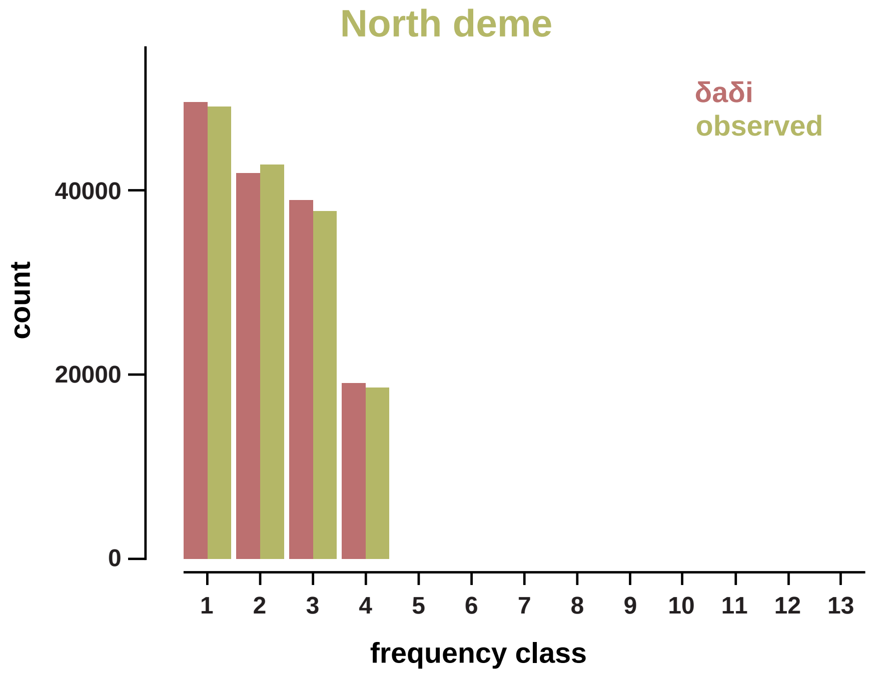

North deme
δaδi
observed
40000
20000
0
count
1
2
3
4
5
6
7
8
9
10
11
12
13
frequency class

c)

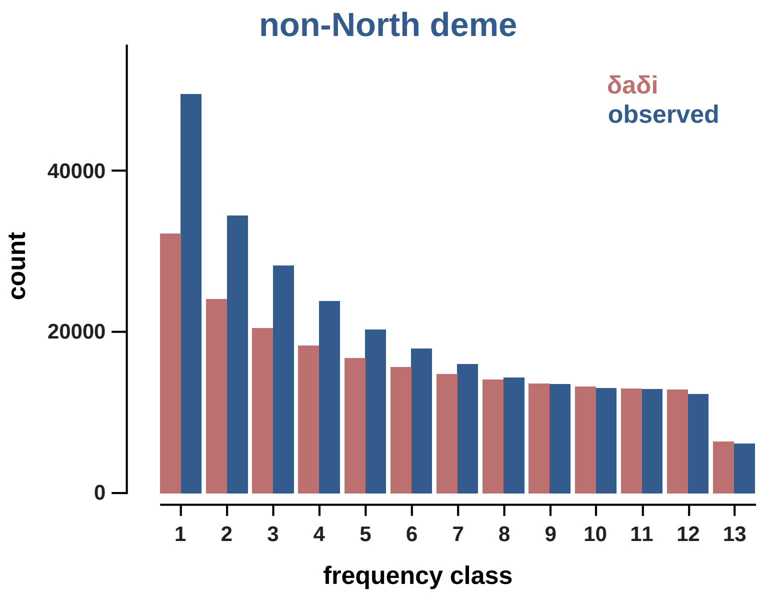


d)

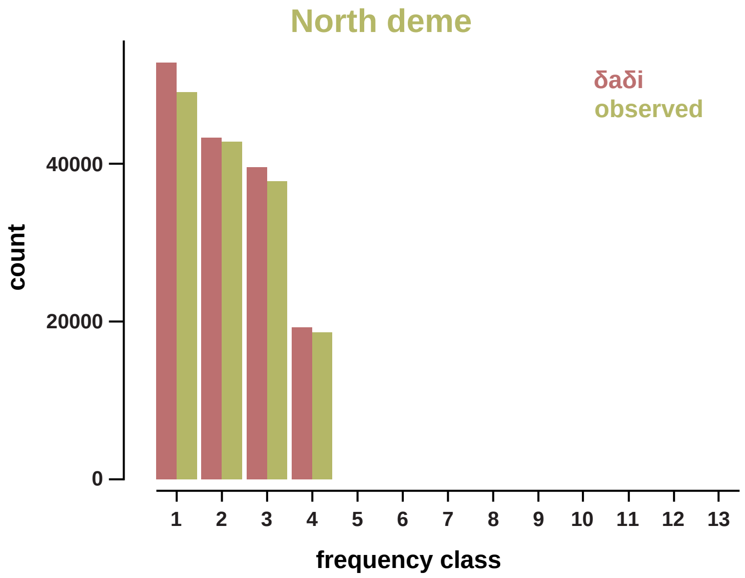


**Supplementary Figure 8**. Comparisons between the observed folded site frequency spectra (SFS) to the calculated SFS generated by *δaδi* for the non-North deme (panel a) and North deme (panel b), and to the mean folded SFS generated by msprime using the estimated model from *δaδi* for the non-North deme (panel c) and North deme (panel d).

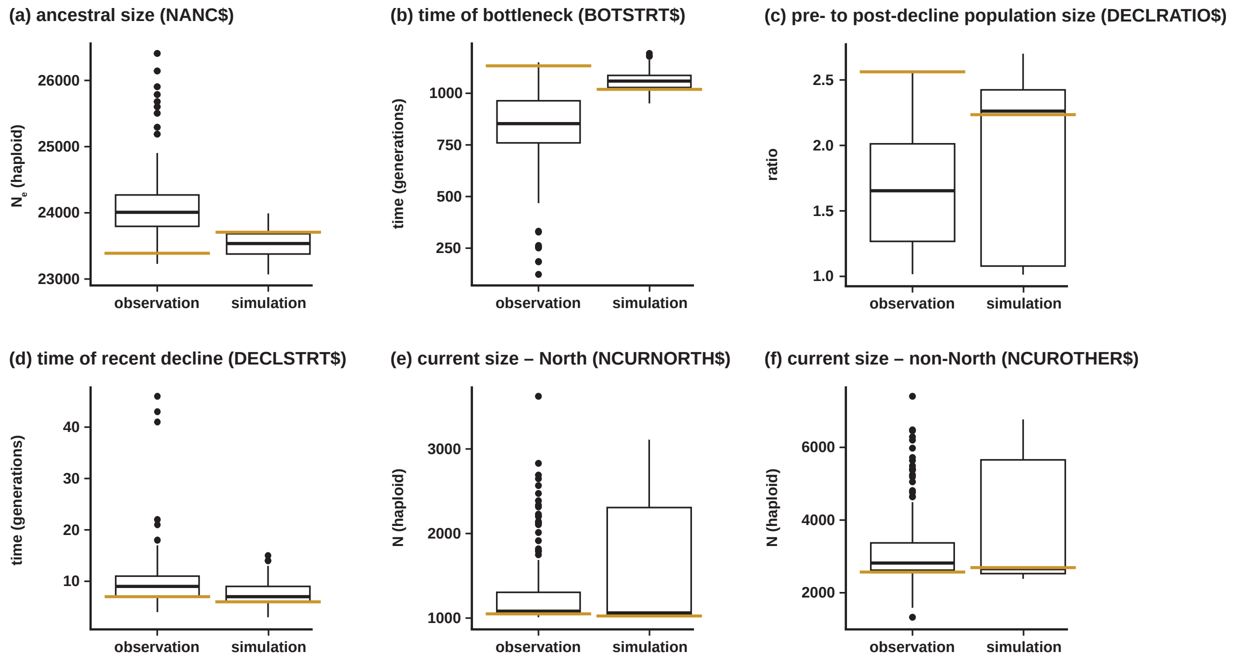


**Supplementary Figure 9**. Box and whisker plots of the range of estimated parameters, comparing estimates obtained from the observed empirical data to the best parameter estimations based on 100 simulated site frequency spectra (SFS), for: (a) the ancestral population size, (b) the timing (in generations) of the population bottleneck, (c) the ratio of the ancestral population size to the post-decline population size, (d) the initiation of the recent population decline, (e) the current size of the North population, and (f) the current size of the non-North population. The orange horizontal line extending past the width of each box marks the parameter value for the best-fitting model.

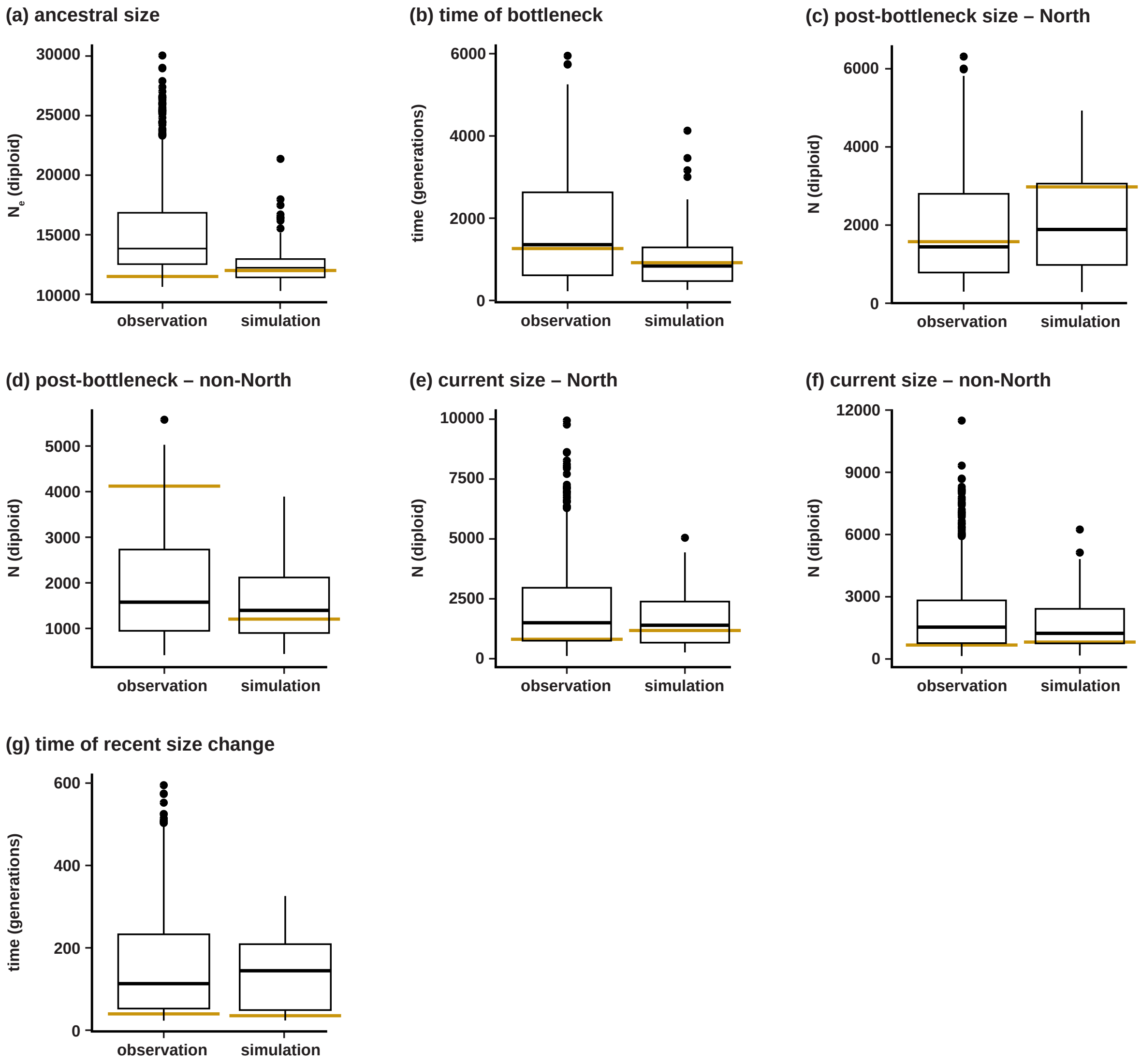


**Supplementary Figure 10**. Box and whisker plots of the range of estimated parameters using *δaδi,* comparing that estimated from the observed empirical data to the best parameter estimations using 100 site frequency spectra (SFS) simulated with msprime, for: (a) ancestral, diploid population size, (b) time (generations) since initial bottleneck, (c) post-bottleneck diploid population size in the North deme, (d) post-bottleneck diploid population size in the non-North deme, (e) current, diploid population size in the North deme, and (f) current, diploid population size in the non-North deme, (g) time (generations) since recent population size change began. The orange horizontal line extending past the width of each box marks the parameter value for the best-fitting model.
